# Toward a probabilistic definition of chromatin accessible regions at the single-cell level

**DOI:** 10.64898/2026.05.01.722232

**Authors:** Elena Sanchez-Escabias, Daniel Rico, Jose C. Reyes

## Abstract

Understanding cis-regulatory elements (CREs) at the single cell level is fundamental to deciphering transcriptional changes during development, cell differentiation, and homeostasis. Recent studies have shown that arbitrary peak-calling thresholds complicate data interpretation and cross-study comparisons. Furthermore, due to the inherent sparsity of single-nuclei ATAC-seq (snATAC-seq) data, distinguishing between truly inaccessible regions and technical dropouts remains challenging. Our analysis of snATAC-seq experiments performed in a well-established cell line suggests that the dichotomy between accessible (open) or inaccessible (close) CREs is misleading. Thousands of accessible regions are present in a very small fraction of cells of the population but they are repeatedly identified, suggesting that they have a low accessibility or are only transiently accessible. However, depending on the detection threshold selected they could be considered as either genuine CREs or noise. To resolve this inconsistency, we propose a model where chromatin accessibility is treated as a continuum, defined by a probability of accessibility (*Pa*) for each accessible region across cell types and conditions. Through computational simulations, we demonstrate that snATAC-seq results can be explained by a simple “balls into bins” probability model, offering a theoretical framework for calculating *Pa* distributions from any snATAC-seq dataset. Furthermore, we examine how *Pa* distributions shift following activation of the TGFβ signaling pathway. This probabilistic approach removes the reliance on arbitrary thresholds and supports a more quantitative, and dynamic understanding of accessible regions’ function.

## INTRODUCTION

Cis-regulatory elements (CREs) of the genome (typically promoters, enhancers, and insulators) mediate spatiotemporal control of gene expression. Traditional bulk approaches have assumed that CREs are in either an active or an inactive configuration, with the active conformation characterized by the presence of accessible chromatin [1-4]. This implies that, in regions where CREs are accessible in a specific cell type, referred to as accessible regions (ARs), nucleases or transposases can interact with the DNA at a higher rate than in other regions of the genome. It is broadly accepted that the degree of chromatin accessibility is determined by the occupancy and topological organization of nucleosomes together with other chromatin factors that block access to DNA [5-7]. Chromatin accessibility has been traditionally measured by quantifying the susceptibility of chromatin to either enzymatic methylation or cleavage [7]. However, today, the most popularized method is the assay for transposase-accessible chromatin using sequencing (ATAC-seq), which leverages the ability of the Tn5 transposase to insert sequencing adapters into ARs at a higher frequency than in nucleosome-occupied regions [8, 9]. Bulk ATAC-seq experiments have been extensively used to map cis-regulatory regions along the genome in different cell types and conditions [10]. However, the extent to which ATAC-seq can be used as a quantitative method that directly measures DNA accessibility, as well as the molecular configuration of regions with a determined ATAC-seq signal, is far from being clearly understood. Thus, the degree of accessibility can be very different from one region to another, and therefore, estimating whether a CRE is active or inactive based on the degree of accessibility can be challenging. Additionally, does an increase in accessibility imply a higher level of the accessibility of the CRE in all cells of the population, or does it reflect an increase in the number of cells where the CRE is accessible? This type of question could, at least theoretically, be addressed using single-nuclei ATAC-seq (snATAC-seq). However, this technology has intrinsic problems, such as data sparsity: less than 4% of the values of peak × cell matrices contain non-zero values, and the lack of consensus on the thresholds used to define what is a genuine accessible region and what is noise. A consequence of this problematic is the significant variation in the number of ARs reported across different snATAC-seq publications. For example, snATAC-seq experiments have reported as many as 620,386 [11] versus 378,207 [12] ARs in human retina, or 437,311 [13] versus 214,890 [14] ARs in kidney, which may be also dependent on the number of cell types identified. Recent works have raised this problem and proposed methods to address data sparsity and CRE identification in snATAC-seq analysis using matrix factorization or multi-factor models [15-17]. While these approaches reduce noise, improve data normalization, and facilitate AR identification, they generally focus on detecting accessible regions rather than quantitatively characterizing individual ARs.

Statistical mechanics is a physical-mathematical framework that applies statistical methods and probability theory to explain macroscopic systems in terms of microscopic parameters, which fluctuate around average values and are characterized by probability distributions. Single-cell genomic analysis is ideal for applying statistical mechanics approaches to understand genomic regulation [18]. This work aims to establish a theoretical framework for defining the degree of accessibility of ARs through probability values *Pa*_*x*_ (where *x* denotes each AR detected in the genome). To do so, we analyzed snATAC-seq data as if it were a typical probability problem of throwing *m* balls into *n* bins. The results enable us to assign a probability of accessibility to each AR. In this model, it is no longer necessary to be concerned about the certainty of the accessible condition of an AR; instead, its state is defined by a probability, which can change after a regulatory stimulus.

## RESULTS

### How many accessible regions are there in the genome of a cell line?

Bulk ATAC-seq experiments present a large heterogeneity in signal intensities at accessible regions (ARs). A low signal in a peak can indicate either generally low accessibility of the AR across most cells in the population, or the presence of only a small subset of cells in which the AR is accessible. To provide a more quantitative description of the nature of ARs, we performed a scMultiome experiment (snATAC-seq + snRNA-seq from the same nuclei) on normal mouse mammary gland epithelial cells (NMUMG) grown under standard conditions (see methods). We then analyzed the snATAC-seq dataset to assess chromatin accessibility in these cells. First, we performed a peak-calling (MACS2 software [19]) using 3925 cells that provided 236,128 ARs (see Methods). Two recent publications have demonstrated the convenience of counting fragments instead of reads to preserve quantitative regulatory information in snATAC-seq experiments [20, 21]. Consequently, we generated an AR × cell matrix with the raw number of fragments using Seurat (v4.3.0.1) and Signac (v1.11.0) software [22, 23]. We found a total of 51,586,278 fragments in the 236,128 ARs analyzed. We did not filter by any minimum threshold in the number of cells to define the ARs. Figure 1A shows the frequency distribution of the number of cells per AR. Interestingly, the mode of this distribution is 22, and the median 46, indicating that most of the detected ARs are only accessible in a small number of cells. In fact, approximately 82% of the ARs were detected only in fewer than 5% of the analyzed cells (Figure 1A).

**Figure 1.**
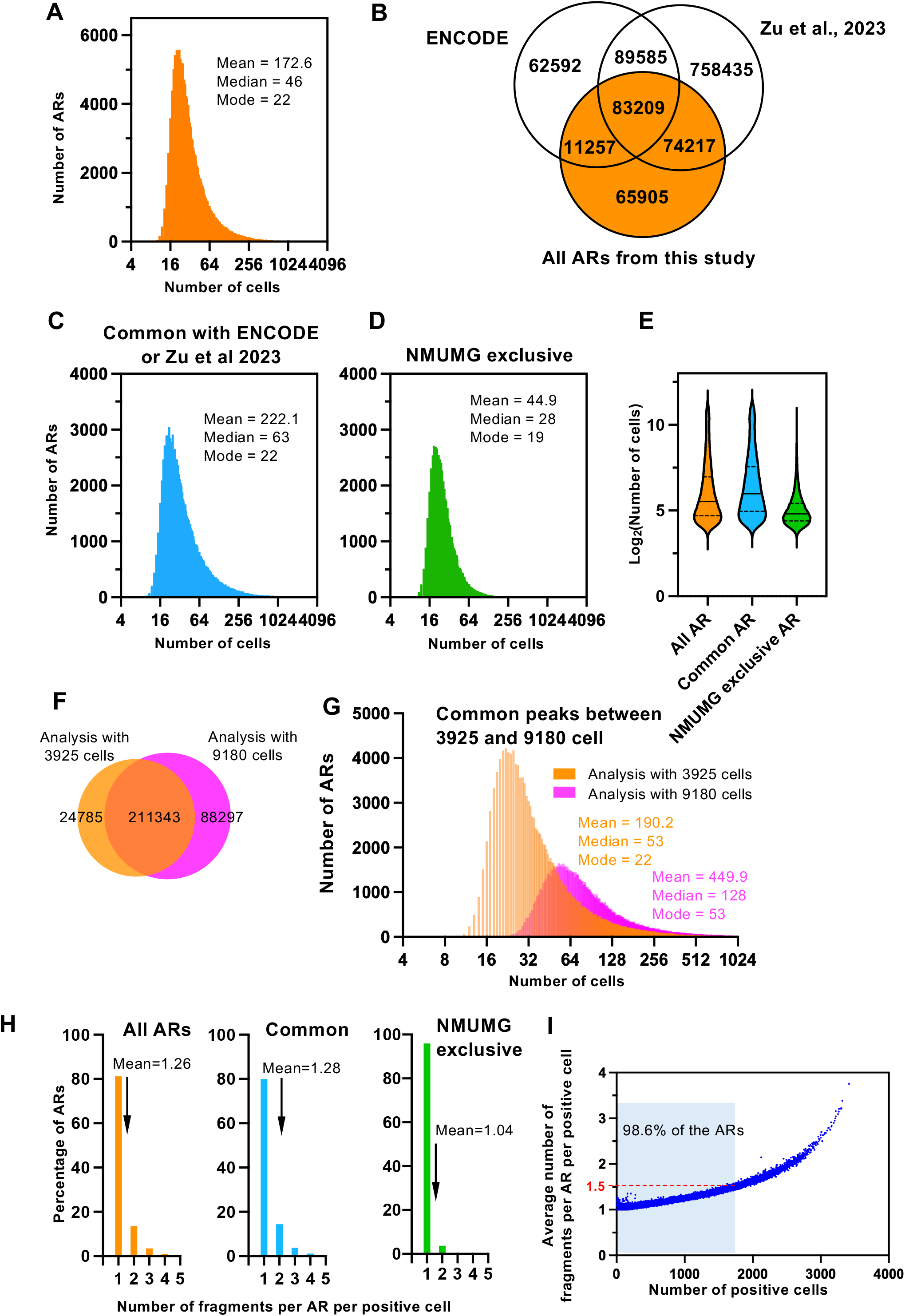
Number of accessible regions in the NMUMG cell line. Accessible regions were detected by snATAC-seq using 3,925 NMUMG cells. **A**. Frequency distributions of the number of positive cells (cells with at least one fragment for the AR) for each AR detected in this study. **B**. Overlapping between ARs detected in this study and mouse ARs previously described at the ENCODE mouse Registry of Candidate cis-Regulatory Element [24] and the candidate CREs from mouse adult brain [25]. **C**. Frequency distributions of the number of positive cells for each common AR between this study and those of ENCODE [24] or mouse adult brain [25] (Common ARs). **D**. Frequency distributions of the number of positive cells for each AR detected in this study but not in ENCODE [24] or mouse adult brain [25] (NMUMG-exclusive ARs). **E**. Violin plot showing the distributions of the number of positive cells per AR for all, common and NMUMG-exclusive ARs. **F**. Comparison between the ARs detected by snATAC-seq using 3925 cells or 9,180 cells. **G**. Frequency distributions of the number of positive cells for each AR, detected using 3,925 cells or 9,180 cells. **A, C, D, G**. Please note that the x-axis is in log_2_ scale. **H**. Distribution of the number of fragments per AR per positive cell for all ARs detected in this study (left), for the common ARs (center) or for the NMUMG-exclusive ARs (right). **I**. Scatter plot showing correlation between the number of positive cells for a specific AR and the average number of fragments per AR per positive cell. 98.6% of the 236,128 NMUMG ARs are located in the shaded area.

Given that the total number of proposed CREs in the mouse or human genomes is around one million [24, 25], it may seem surprising that a single cell line, NMUMG, has 236,128 ARs, raising the question of whether ARs detected in only a few cells are biologically relevant or noise. To address this question, we have compared our list of ARs with two lists of known proposed CREs in the mouse genome: the ENCODE registry of candidate cis-regulatory elements (339,815 ARs) [24] and a recently published list of candidate CREs from the mouse adult brain, which includes 1,053,811 ARs across 1,482 distinct brain cell populations [25]. As shown in Figure 1B, more than 71% of the ARs identified in this study overlapped with ARs previously identified in one of these two works, suggesting that they represent true CREs, even though many were accessible in only a very small fraction of the NMUMG cell population. Furthermore, when we analyzed only the ARs previously identified in these two works, the distribution of the number of cells in which a specific AR was detected resembled that obtained using the full list of 236,128 ARs detected in NMUMG cells (Figure 1C, 1E). In the case of NMUMG-exclusive ARs, while the mode was similar to that of the other two distributions (19 versus 22 cells), the median and the mean were clearly smaller (Figure 1D and 1E), indicating that, on average, these regions have a lower tendency to be accessible. However, since most of these ARs are detected in a number of cells similar to others previously identified, there is no reason to rule out these putative CREs in NMUMG cells. Additionally, further analysis of NMUMG-exclusive ARs revealed that they were less enriched in TSS, presented a lower GC content and had a shorter width (Supplementary Figures S1A, S1B and S1C), suggesting they might be mainly associated with NMUMG-specific enhancers.

These data lead us to hypothesize that many CREs have an intrinsic probability of being accessible in nearly any cell type, possibly due to the low tendency of the underlying DNA sequence to position nucleosomes [26-29]. This nucleosome refractory property may lead to transient and sporadic destabilizations of the CRE nucleosomes, creating windows of opportunities for the Tn5 transposase to integrate. If this hypothesis were true, we would expect to find a considerably larger number of ARs by increasing the number of analyzed cells. We then performed peak-calling using data from 9,180 cells from the same experiment described above, which identified 296,898 ARs (Figure 1F). Most of the ARs detected only in a very small number of cells in our first analysis were now detected again, but now in a larger number of cells, suggesting that they were not just noise, but truly detected ARs (Figure 1G). In addition, 88,297 more ARs were detected, confirming our hypothesis. The newly identified ARs presented a frequency distribution of cells with median 42 and mean 48.7, suggesting that, on average, they were found in a small number of cells of the population. However, many of them overlapped with ARs previously identified (Supplementary Figures S1D and S1E). The new peak-calling also failed to identify 24,785 ARs previously detected, which is likely related to the fact that the MACS2 algorithm considers surrounding reads to define accessible regions. Most of these ARs were present in a small number of cells (Supplementary Figure S1F). The strong dependence of the number of identified ARs on the number of cells analyzed highlights the weakness of the identification method and raises doubts about the accuracy of the detected ARs.

Analysis of the fragment frequency in the 236,128 ARs dataset revealed an almost binary (0, 1) distribution, as previously reported in several studies (see for example [8, 20, 21]) (Supplementary Figures S1G, S1H and S1I). We defined as “detected ARs” as those ARs identified by MACS2 for which at least one fragment has been sequenced. A cell is considered “positive” for a specific AR if it contains at least one fragment corresponding to that AR. We found an average of 1.26 fragments per AR per positive cell (Figure 1H). This value was very similar (1.28) for previously identified ARs in [24, 25] and slightly lower (1.04) for the NMUMG-exclusive ARs (Figure 1H). Additionally, we observed a positive correlation between the number of positive cells for an AR and the average number of fragments per positive cell for that AR (Figure 1I). Notably, 98.6% of the detected ARs had a very low average number of fragments per positive cell: between 1 and 1.5. We also observed a positive correlation between the average number of fragments per AR per positive cell and the size of the AR (R^2^=0.45) (Supplementary Figure S1J). The average size of the snATAC-seq sequenced fragments was approximately 164 bp (Supplementary Figure S1K), while the average size of the ARs was 620 bp (ranging from 199 to 5,518 bp). Therefore, and taking into account the two chromosomal copies of a genomic position in a diploid cell, we reason that more than one fragment per AR per positive cell could theoretically be expected. However, due to the physicochemical and kinetic properties of the Tn5 enzyme, the possibility that both genomic positions may not be simultaneously accessible, the crowded nuclear environment, and likely the dilution of the active transposase, only one fragment per positive cell is typically released from most ARs [8].

We conclude the following from these data: First, by analyzing a single cell line, we can detect as ARs approximately one quarter of all known mouse putative CREs. However, while some ARs are detected in only a few cells, others are found in most cells, indicating substantial heterogeneity and suggesting that some ARs may be only transiently accessible. Second, at the single-cell level, both ARs detected in many cells and those detected in only a few are typically represented by a single fragment per AR per positive cell. Therefore, we consider that ARs appearing in only a few cells are not noise, but true accessible regions, at least in those specific cells and at that particular moment. Third, the considerable variation in accessibility makes it misleading to talk about the number of “open” (accessible) and “close” (inaccessible) CREs in a cell type, as a binary property. Instead, we propose to consider chromatin accessibility as a continuum where there are no distinct categories of “open” CREs and “close” CREs, which would require arbitrary thresholds in snATAC-seq experiments. Rather, we suggest modeling a distribution of accessibility probabilities for each accessible regions in every cell type and condition.

### snATAC-seq experiments can be simulated using the “balls into bins” probability problem

Next, we aimed to develop a simple theoretical framework to assign a probability of accessibility to each AR. The observation that most of the ARs are represented by 0, 1, or 2 fragments is reminiscent of the classical probability problem known as “balls into bins”. This problem involves distributing *m* balls randomly into *n* bins of identical size (Figure 2A). In this type of problems, when *m* << *n*, the initial balls almost always land in empty bins, resulting in an average number of approximately one ball per occupied bin. Specifically, the probability that ball_*i*_ (with *I* = 1, 2, …*m*) falls into bin_*j*_ (with *j* = 1, 2, …*n*) is 1/*n*, making the likelihood of two balls landing in the same bin very low (1/*n*^2^). As the number of balls (*m*) increases, the probability of landing in an already occupied bin also rises.

**Figure 2.**
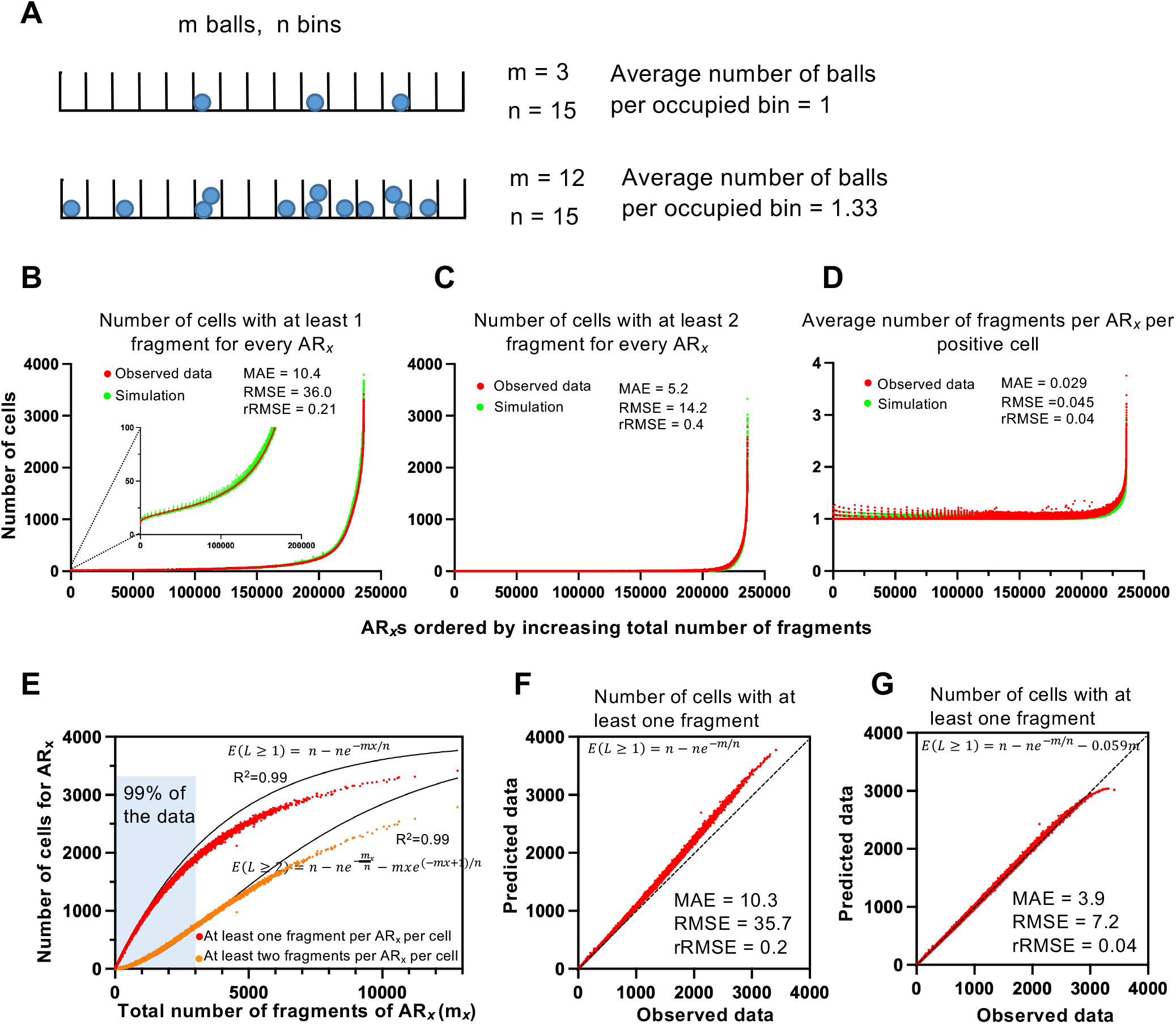
A. snATAC-seq experiments can be simulated using the “balls into bins” probability problem. **A**. Diagram of a balls into bins problem. In the first row 3 balls (*m* = 3) are thrown randomly into an array of *15* bins (*n* = 15) of identical size. For *m* << *n* the number of balls per bin use to be 0 or 1. In the second row *m* = 12 and *n* = 15, now *m* ≈ *n*, and some bins appears with 2 balls. **B-D**. Simulation of the snATAC-seq experiments using a balls into bins strategy were the number of balls (*m*_*x*_) corresponds to the fragments detected for AR_*x*_, and the number of bins (*n*) corresponds to the number of analyzed cells (*n* = 3925). The simulation was iterated 236,128 folds, corresponding to the number of identified AR_*x*_ (with *x* = 1, 2….236,128). Observed values (red) and the simulated values (green) of the number of positive cells for AR_x_ (cells with at least one fragment for AR_x_) (B), the number of cells with more than one fragment for AR_x_ (C) and the average number of fragments per AR_x_ per positive cell (D), are shown. **E** Scatter plot of observed data of the number of cells with at least one fragment per AR_*x*_ (red) or with at least two fragments per AR_*x*_ (orange) versus the total number of fragments of AR_*x*_. Equations 7 and 8 were fitted to the data. Coefficient of determination (R^2^) is provided. **F, G**. Observed versus predicted plots. Predicted values were estimated using the indicated equation, where *n* = 3925 and *m* is the total number of fragments for every AR_*x*_. **B-D, F, G**. Mean absolute error (MAE), root mean square error (RMSE) and relative RMSE (rRMSE).

We hypothesized that the accessibility of a given AR (AR_*x*_ where *x* = 1, 2, …, 236,128) could be modeled using this framework, assuming that *m*_*x*_ represents the total number of fragments (balls) detected for AR_*x*_, and *n* corresponds to the number of analyzed cells (bins). To test this hypothesis, we conducted a “balls into bins” simulation using the observed fragment count for each AR_*x*_ as *m*_*x*_ and the total number of analyzed cells (n = 3,925) as the bins. The simulation was repeated for each identified AR_*x*_. Figures 2B, 2C and 2D show observed experimental values (red) and simulated values (green) for three cases: Figure 2B presents the number of cells containing at least one fragment for AR_x_ (positive cells), Figure 2C shows the number of cells containing more than one fragment for AR_x_, and Figure 2D reflects the average number of fragments per AR_x_ per positive cell. The low mean absolute error (MAE), root mean square error (RMSE), and relative RMSE (rRMSE) suggest that the simulated data closely resemble the experimental data.

The “balls into bins” problem can be analyzed using either the Binomial or the Poisson distributions (see methods). Here, we derive expressions for key probabilities using the Binomial approximation, while the corresponding equations for the Poisson distribution are presented in Material and Methods. Let *L* be a discrete random variable with a Binomial distribution showing the number of balls in the *j-*th bin. If the probability that ball_*i*_ falls into bin_*j*_ is 1/*n*, then, the probability mass function for *L* is

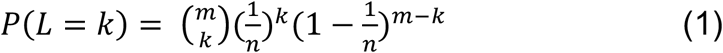

where *k* is the number of balls in the bin.

Then, the probability of having an empty bin is:

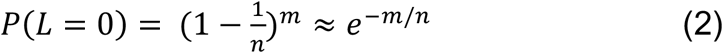

The probability of having at least one ball is:

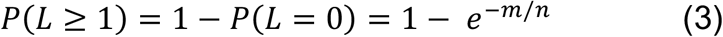

and the probability of having at least two balls is:

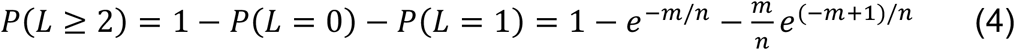

The corresponding expectations, applying the linearity of the expectation property, are

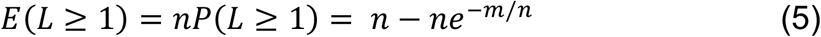

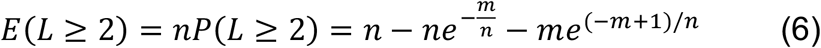

Next, we tested whether equation 5 and 6 fit the observed values of cells with at least one fragment for AR_*x*_ and the observed values of cells with at least two fragments for AR_*x*_, respectively (Figure 2E, 2F). Approximately 99% of the observed AR_*x*_ data were well fitted by equations 5 (R^2^ = 0.99) and 6 (R^2^ = 0.99).

However, for AR_*x*_s with a large total number of fragments (m_*x*_ > 2,000), the observed values of cells were slightly lower than expected (Figure 2E, 2F and Supplementary Figure S2A). This suggests that for highly accessible ARs, the transposase tends to insert more frequently than predicted by the model. In the analogy of the “balls into bins” problem, balls would have a small preference for already occupied bins. To improve the fit, we introduced a factor dependent on the total number of fragments detected, which strongly enhanced the fit for the number of cells with at least one fragment for each AR_*x*_ (equation 7) (compare Figure 2F and 2G, and Supplementary Figures S2A and S2B).

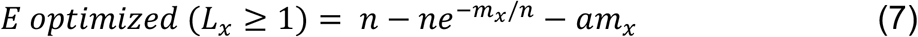

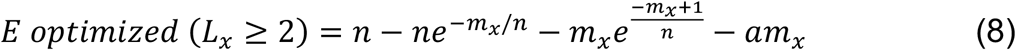

Where *a* is a hyperparameter that we set empirically (see methods).

Next, we assume that equation 7 represents the theoretical expectation of finding cells with AR_*x*_ (i.e., finding AR_*x*_ actually accessible). Therefore, since *E*(*L*_*x*_ ≥ 1) = *nP*(*L*_*x*_ ≥ 1) for the Binomial and Poisson distributions, then, *E*(*L*_*x*_ ≥ 1)/*n* is the probability that a given AR_*x*_ is actually accessible at the cellular level in our experiment. We refer to this parameter as the Probability of accessibility of AR_*x*_ (*Pa*_*x*_), where *x* = 1, 2, … 278,380.

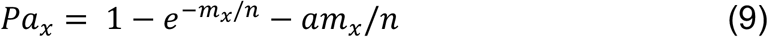

We have calculated *Pa*_*x*_ for all the 278,380 AR_*x*_s identified in our snATAC-seq experiment (Figure 3A), with values ranging from 0.0022 to 0.774. This indicates that those AR_*x*_s not detected have a *Pa*_*x*_ < 0.0022 in our experiment. Figure 3B shows the distribution of *Pa*_*x*_ values observed in our experiment, with only 10.3 % of the AR_x_s having *Pa*_*x*_ > 0.1. *Pa*_*x*_ is slightly positively correlated to the AR_*x*_ width (Supplementary Figure S2C). Very similar results were obtained when using the data obtained from 9,180 cells (Supplementary Figures S2D, S2E, S2F, S2G), indicating that the parameter *Pa*_*x*_ is not sensitive to the number of cells used in the analysis. However, *Pa*_*x*_ was sensitive to the sequencing depth as evidenced by the decrease of *Pa*_*x*_ values when the average number of AR fragments per cell was decreased by downsampling (Supplementary Figure S2H).

**Figure 3.**
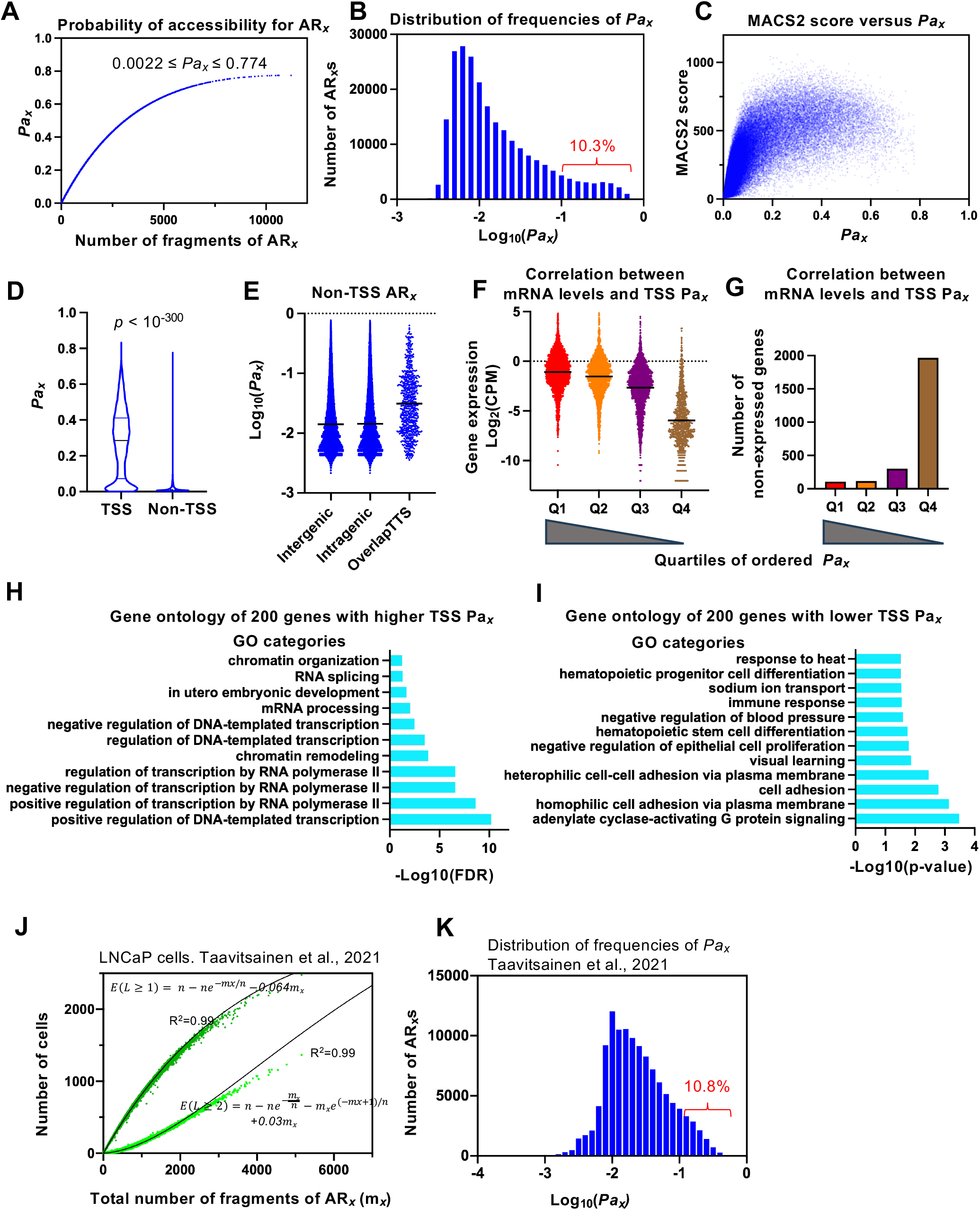
Characterization of the Probability of accessibility (*Pa*_*x*_) parameter. **A**. Scatter plot showing how Pa_*x*_ depends on the total number of fragments detected for AR_*x*_. **B**. Distribution of frequencies of *Pa*_*x*_ for all AR_*x*_ detected in this study. Data are binned into intervals of 0.1 units. Bars are represented in the center of the interval. **C**. Scatter plot showing the relationship between MACS2 score and *Pa*_*x*_. Each dot represents an AR. **D**. Distribution of *Pa*_*x*_ values in TSS and non-TSS ARs. **E**. Distribution of *Pa*_*x*_ values in non-TSS ARs from different genomic locations. **F, G**. TSS were ordered from higher to lower *Pa*_*x*_ values and divided into four quartiles. mRNA levels (CPM, from scRNA-seq) of the corresponding genes (F) and the number of non-expressed genes (G) in each quartile are shown. **F**. Only values of the expressed genes are displayed. **H, I**. Biological process GO categories enriched among the 200 genes with higher (H) or lower (I) TSS *Pa*_*x*_ values. **J**. Scatter plot showing data of accessible regions from LNCaP cells [33]. The number of cells with at least one fragment per AR_*x*_ (green) or with at least two fragments per AR_*x*_ (bright green) versus the total number of fragments of AR_*x*_ are plotted. Data were fitted to equations 7 and 8. Coefficient of determination (R^2^) is provided. **K**. Distribution of frequencies of *Pa*_*x*_ for all AR_*x*_ detected in LNCaP cells. Data are binned into intervals of 0.1 units.

The *MACS2 score* is a quantitative metric assigned by the MACS2 peak-calling algorithm that reflects the statistical confidence of chromatin accessibility or binding enrichment at a genomic region [19]. Specifically, MACS2 models the background noise using a dynamic Poisson distribution and estimates the significance of the observed read enrichment compared to the expected background. Therefore, we investigated the relationship between the MACS2 score and *Pa*_*x*_. As shown in Figure 3C, the two parameters correlate positively for regions with low MACS2 scores; however, for MACS2 values greater than approximately 250, *Pa*_*x*_ exhibits substantial divergence. To illustrate this divergence, we selected two genomic regions containing several ARs and compared their respective *Pa*_*x*_ and MACS2 scores. As shown in Supplementary Figure S3A, AR_b_ and AR_g_ both have a MACS2 score of 508, yet their *Pa*_*x*_ values differ markedly (*Pa*_*b*_ = 0.014 and *Pa*_*g*_ = 0.468). This indicates that, despite similar enrichment significance, the probability of observing accessibility across cells is much lower for AR_b_ than for AR_g_. In other words, very few cells exhibit accessibility at AR_b_, whereas AR_g_ is accessible in a larger fraction of the cell population. Therefore, we concluded that MACS2 score and *Pa*_*x*_ provide different information.

*Pa*_*x*_ values were much higher for regions annotated as TSS compared to non-TSS regions (Figure 3D). Interestingly, among the non-TSS regions, *Pa*_*x*_ was higher in ARs overlapping transcription termination sites (Figure 3E), which may be related to the expression of antisense lncRNAs involved in transcription termination from enhancers at the 3’end of the genes [30, 31]. We observed a modest positive correlation between *Pa*_*x*_ for TSS and mRNA levels (snRNA-seq data) from the same scMultiome experiment (R^2^ = 0,27) (Figure 3F and Supplementary Figure S3B). In fact, expression of approximately 67% of the genes with a TSS *Pa*_*x*_ below 0.086 was not detected in the snRNA-seq from the scMultiome experiment (Figure 3G). Interestingly, TSS from genes with very high mRNA levels, such as ribosomal protein genes, did not display the highest *Pa*_*x*_ values (Supplementary Figure S3C). Next, we performed a functional analysis of genes with higher and lower *Pa*_*x*_. We observed a strong overrepresentation (25%) of genes from chromosome 19 among the genes whose TSS exhibited the top 200 *Pa*_*x*_ values (Supplementary Figure S3D), likely due to the trisomy of this chromosome in NMUMG cells [32]. To eliminate potential bias, we excluded genes from chromosome 19 for this analysis. Notably, genes with the highest 200 TSS *Pa*_*x*_ values were enriched in Gene Ontology categories related to transcriptional regulation, chromatin organization and RNA splicing (Figure 3H). In contrast, genes with the lowest 200 TSS *Pa*_*x*_ values were enriched in tissue specific categories and cell adhesion (Figure 3I), and their mRNA expression was either undetected or very low.

Next, we verified that the “balls into bins” approximation can be applied to other set of single-cell accessibility data. Equations 7 and 8 fit the data from snATAC-seq experiment on the human LNCaP cell line [33] perfectly (R^2^ = 0.99) (Figure 3J). We then calculated *Pa*_*x*_ for each detected AR_*x*_, obtaining a *Pa*_*x*_ distribution similar to that observed for NMUMG cells (Figure 3K and Supplementary Figure S2I).

We have made publicly available our script to compute *Pa*_*x*_ (https://github.com/elesanesc/scPaX) and it can be incorporated easily into standard pipelines.

### Signal transduction pathways change *Pa*_*x*_ values

TGFβ cytokine signaling is transduced through the phosphorylation and activation of receptor-regulated (R-) SMAD proteins (SMAD3, among others). R-SMAD factors oligomerize with SMAD4 and translocate to the nucleus, where they bind chromatin [34]. The activated SMAD3/SMAD4 complexes are typical signal-driven TFs, predominantly binding enhancers in association with other transcription factors [2, 35]. Previously, we showed that TGFβ leads to an increase in the bulk ATAC-seq signal in numerous CREs that were already accessible before TGFβ treatment in NMUMG cells [36]. This observation prompted the question of whether the increase in accessibility was due to enhanced CRE opening in all cells, or whether it reflected an increase in the number of cells with accessible CREs. We speculated that a snATAC-seq experiment may help us to decide between these two possibilities. To address this, we compared snATAC-seq data from TGFβ-treated cells with data from untreated control cells. To ensure a fair comparison of the different parameters, including *Pa*_*x*_, we normalized the average number of fragments per cell by downsampling, to match that of the sample with the lowest sequencing depth. Then, we performed peak calling in this new sample using MACS2 and generated a consensus list of 350,123 ARs.

After differential accessibility analysis using FindMarkers() function from the Seurat and Signac packages [22, 23], we found that TGFβ increased the accessibility of 111,359 AR_x_s (32% of the AR_x_s) (p-value< 0.05 and Log_2_FC >0.5), and decreased the accessibility of 64,077 AR_x_s (18%) (p-value< 0.05 and Log_2_FC < 0.5). For TGFβ-increased AR_x_s, the number of cells where some AR_x_s were identified increased between 1.5- and approximately 60-fold in TGFβ treated cells compared to the control cells (Figure 4A, 4B and 4D). In contrast, the number of fragments per cell for detected AR_x_ changed between 0.5- and 1.45-fold (Figure 4C, 4D and Supplementary Figure S4A). Interestingly, the largest increases in the number of positive cells following TGFβ treatment occurred in AR_x_s that were initially detected in a small number of cells (Supplementary Figure S4B and S4C). In contrast, the largest increases in the number of fragments occurred mostly at AR_x_s detected in a large number of cells (Supplementary Figure S4D and S4E). Despite the substantial increase in the number of cells with accessible regions for many AR_x_s, the average number of fragments per AR_*x*_ per positive cell remained close to 1 in most cases (Figure 4C, 4D). Surprisingly, for some AR_x_s that presented a strong increase in the number of cells, a slight decrease in the fold change of the average number of fragments per cell was observed (data highlighted in light blue in Figure 4D). Equation 7 fits almost perfectly the new TGFβ data, with minimal changes in parameter *a*, respect to the control data (0.042 ± 0.00015 versus 0.028 ± 0.00010) (Figure 4E). This suggests that TGFβ does not drastically alter the snATAC-seq experimental conditions and can therefore be modeled using the “balls into bins” paradigm. Consistently, a “balls into bins” simulation, where the number of balls was increased in proportion to the total number of fragments per each AR_x_ due to TGFβ, successfully recapitulated similar fold changes in both the number of positive cells for each AR_x_ and the average number of fragments per AR_x_ per positive cell (Figures 4F, 4G). Notably, the simulation also reproduced the observed decrease in the fold change of the average number of fragments per cell for some AR_*x*_s that presented an increase in the number of cells (compare light blue data in Figure 4D with Figure 4F). A closer inspection of this phenomenon revealed that it occurs in AR_*x*_s detected in only a very small number of cells, suggesting that it results from the random, sporadic occurrence of two fragments per AR_*x*_ in some cells under control conditions. Taken together, these results suggest that the number of cells and the number of fragments per AR_*x*_ are linked variables, driven by the Poisson-like nature of fragments occurrence at AR_x_s. Accordingly, snATAC-seq experiments cannot distinguish whether an observed increase in snATAC-seq signal reflects a higher accessibility of the CRE in all the cells of the population or simply an increase in the number of cells with the accessible CREs. To address this limitation, we propose using *Pa*_*x*_, which avoids this dichotomy that we have shown to be meaningless. Therefore, we computed *Pa*_*x*_ after TGFβ treatment for the 111359 and 64,077 AR_x_s that increased or decreased, respectively, in accessibility. Figure 4H shows the distribution of *Pa*_*x*_ values of TGFβ-increased AR_x_s before and after treatment. Similar results were observed for AR_x_s that decreased in accessibility (Supplementary Figures S4F to S4K). Interestingly, under control conditions, the average *Pa*_*x*_ of TGFβ-decreased AR_x_s was substantially higher than that of TGFβ-increased AR_x_s (0.071 vs. 0.013). This suggests that TGFβ primarily increases the accessibility of AR_x_s that, on average, are present in a small number of cells under control conditions, while decreasing the accessibility of AR_x_s that are widespread across many cells.

**Figure 4.**
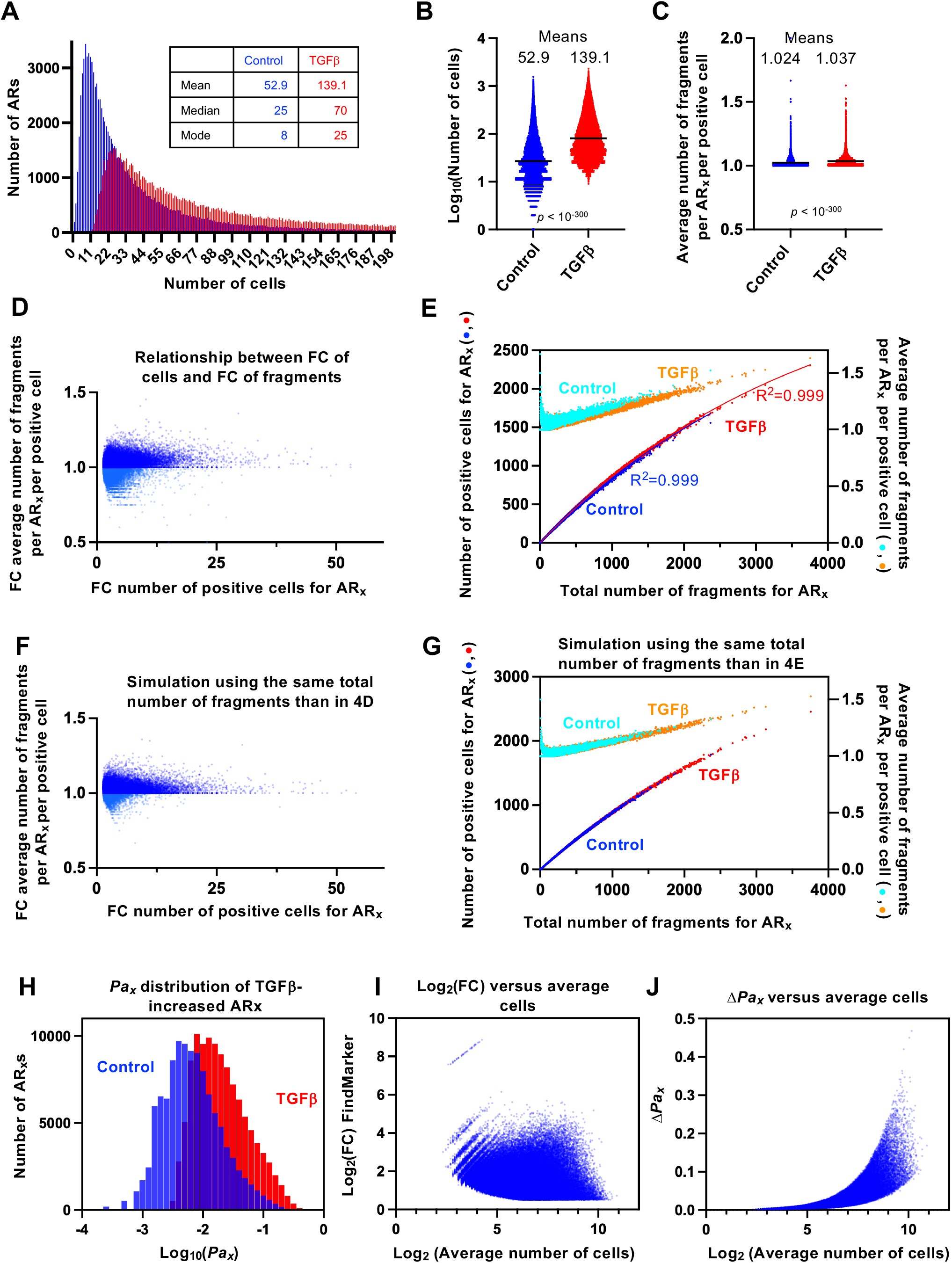
TGFβ alters the probability of accessibility but not the identity of the ARs. A scMultiome experiment of NMUMG cells treated for 24 h with TGFβ was performed and the snATAC-seq results were compared with those of non-treated cells described above. Data displayed in this figure correspond to the 111,359 AR_*x*_s that increased accessibility upon TGFβ treatment (TGFβ-increased AR_*x*_s). Data for TGFβ-decreased AR_*x*_s are shown in Supplementary Figure S4F to S4M. **A**. Frequency distributions of the number of positive cells for the TGFβ-increased AR_*x*_s. Only the first part of the distribution is shown. **B**. Distribution of the number of positive cells in TGFβ-treated and non-treated cells. **C**. Distribution of the average number of fragments per AR_*x*_ per positive cell in TGFβ-treated and non-treated cells. **D**. Relationship between the fold change (FC, TGFβ versus control) of the number of positive cells for AR_*x*_ and the FC of the average number of fragments per AR_*x*_ per positive cell. **E**. Scatter plot of the total number of fragments of AR_*x*_ versus the number of positive cells for AR_*x*_ (left axis). TGFβ (red) and control (blue) data of the same AR_*x*_s are displayed together for comparison. Equation 7 is fitted to the TGFβ data. Coefficient of determination (R^2^) is provided. The total number of fragments of AR_*x*_ versus the average number of fragments per AR_*x*_ per positive cell is displayed in the right axis. TGFβ (orange) and control (light blue) data of the same AR_*x*_s are displayed together for comparison. **F, G**. Same than D and E but displaying simulated data using the balls into bins model. Simulations were performed using the number of fragments identified for TGFβ-increased AR_*x*_ in TGFβ and control conditions as balls. The number of positive cells, the number of fragments per AR_*x*_ per positive cell and FCs (TGFβ versus control) are obtained from the simulation. **H**. Comparison of the distributions of frequencies of *Pa*_*x*_ under control or after TGFβ conditions, for all TGFβ-increased AR_*x*_s. Data are binned into intervals of 0.1 units. **I**. MA plot showing Log_2_FC (from FindMarkers(), Signac) values versus Log_2_(average number of cells) for TGFβ-increased AR_x_. **J**. MA plot showing Δ*Pa*_*x*_ values versus Log_2_(average number of cells) for TGFβ-increased AR_x_.

Fold-change (FC) values alone can be misleading, as their biological relevance depends on the underlying signal magnitude. To obtain a more meaningful measure of accessibility variation in terms of the cell population, we propose using the difference in *Pa*_*x*_ (ΔPa_x_) between conditions instead of or in addition to the snATAC-seq signal FC. As shown in the MA plots of Figure 4I and 4J for TGFβ-increased AR_x_s and in Supplementary Figures S4L and S4M for TGFβ-decreased AR_x_s, Δ*Pa*_*x*_ and Log_2_FC (computed with Signac) provide very different information. While a high Log_2_FC can be associated with changes only in a few cells of the population, a high Δ*Pa*_*x*_ indicates changes that strongly impact in the proportion of cells where AR_x_ can be detected.

Figure 5 shows several examples that demonstrate the usefulness of using *Pa*_*x*_ and Δ*Pa*_*x*_ to gain a more realistic understanding of chromatin accessibility in single-cell experiments. For example, in Figure 5A, peaks b, e and f display similar Log_2_FC (2.7, 2,2, 2.3, respectively). However, reporting Δ*Pax* values reveal a crucial distinction: peak b and e have a Δ*Pa*_*x*_ of 0.177 and 0.098 respectively, indicating detection in a substantial proportion of cells, whereas peak f has a Δ*Pa*_*x*_ of only 0.018, signifying that it remains present in only a small fraction of cells. Thus, Δ*Pa*_*x*_ conveys biologically meaningful information that FC alone obscures. In addition, the TSS of the *Lpin3* gene (AR_b_ in Figure 5A) changed its *Pa* from 0.032 in control conditions to 0.209 after 24 h of TGFβ. While this denotes a strong increase in accessibility, it by no means implies that this region is accessible in all cells of the population, as can be observed in the UMAP in Figure 5B. The fact that *Pa*_*b*_ = 0.209 clearly indicates this fact. Similar cases are the TSS of the *IL11* and the enhancer close to *Hs6st2* genes (Figures 5C, 5D, 5E, 5F). The TSS of the *Rpl28* and *Ube2S* genes had a *Pa* of around 0.32 and 0.59 in control conditions. Even in these cases, the snATAC-seq experiment only detected these regions in a fraction of the cells, making it impossible to experimentally determine whether the non-detection in many cells is a technical impossibility or whether the region is truly inaccessible in those cells, at that moment. Our proposal bypasses this issue, and the state of the region is defined by the determined *Pa*_*x*_.

**Figure 5.**
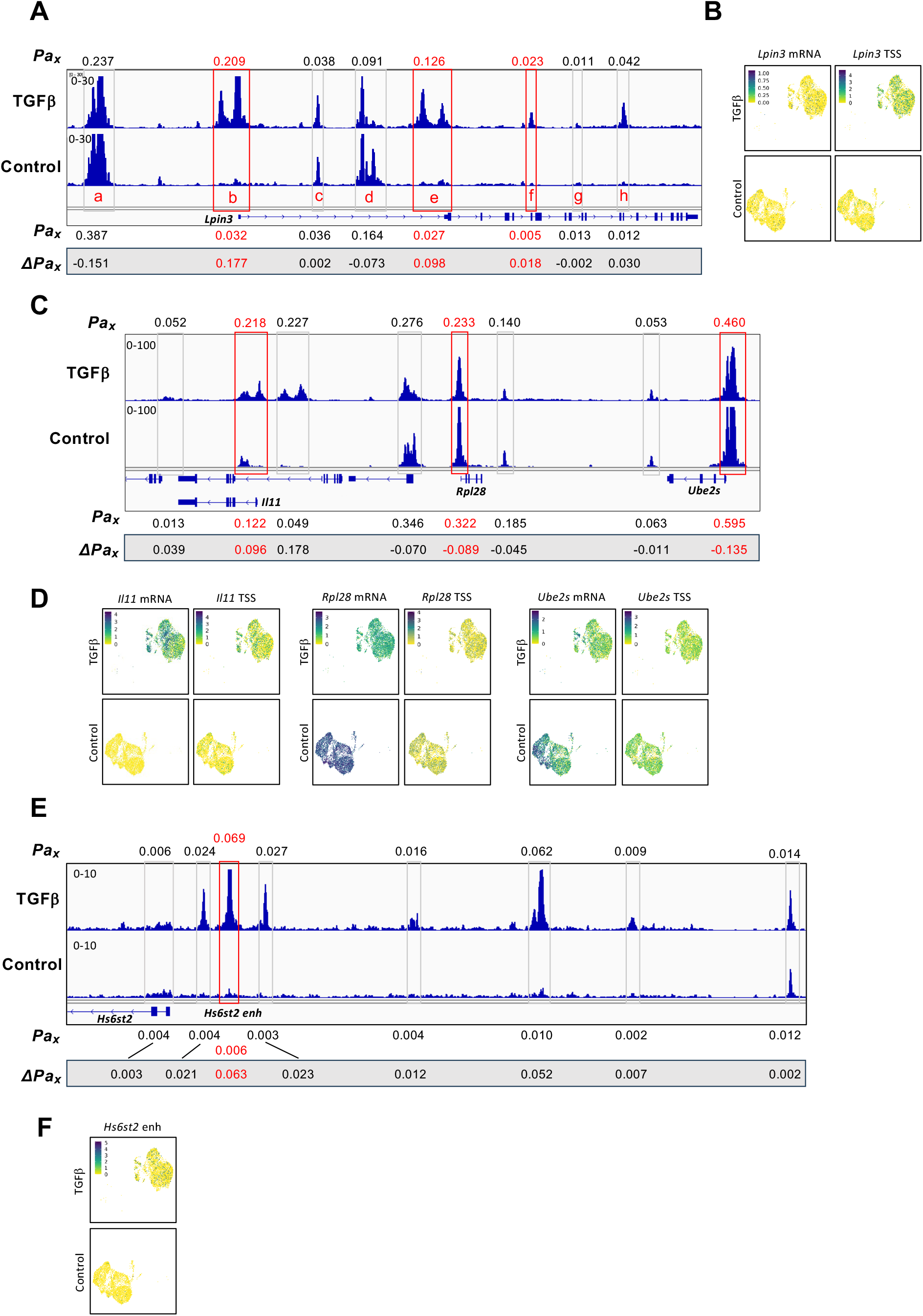
*Pa*_*x*_ provides a more realistic and quantitative description of accessibility. **A, C, and E**. Genome browser snapshot of three genomic regions showing pseudobulk snATAC-seq peaks (ARs) of NMUMG cells under control conditions or 24 hours after TGFβ addition. *Pa*_*x*_ and Δ*Pa*_*x*_ values of the squared ARs are shown. **A**. Letters in red designate ARs referred to in the text. **B, D, F**. UMAPs from the integration of snATAC-seq and scRNA-seq data (from the same scMultiome experiment) from NMUMG cells under control conditions or 24 hours after TGFβ addition, colored by the indicated gene marker expression (scRNA-seq) or AR signal (snATAC-seq).

## DISCUSSION

It is commonly assumed that the set of accessible CREs in the genome defines different cell types. However, due to inherent cellular heterogeneity, the sparsity of snATAC-seq data, the vast differences in accessibility between elements, and the different signal threshold used across studies, determining whether a CRE is or is not accessible in a given cell type is often challenging. In this work, we propose a paradigm shift in defining accessible regions in a cell type by adopting a probabilistic framework. Instead of a binary classification, we define each AR by a probability of being accessible in the cell population (*Pa*_*x*_) and a cell type by a distinct *Pa*_*x*_ distribution. This quantitative and more dynamic approach aligns well with the current understanding of cell types in both normal and pathological processes as a continuum of states [37-41].

The Poisson distribution is broadly used in genomics. Thus, the widely used MACS software assumes that, in the absence of a “true peak,” sequenced reads are distributed approximately randomly across the genome according to a Poisson distribution [19]. However, MACS does not provide the probability of detecting accessible regions from a specific region across a cell population; rather, it identifies genomic regions that contain more reads than would be expected by chance. Recent publications have also presented very robust computational frameworks that estimate the probability of chromatin accessibility based on the Poisson or the Bernoulli distributions [15, 17, 21, 42]. Nevertheless, these methods neither use the determined probability values as intrinsic qualifiers that characterize each AR, nor use probability of accessibility values for downstream analysis, such as comparative quantitative analyses across experimental conditions.

Our data suggest that most accessible regions detected in a snATAC-seq experiment are detected in only a small fraction of cells (82% of the ARs are detected in fewer than 5% of the cells) and therefore have a low probability of detection. However, many of these regions correspond to CREs, mostly enhancers, that are strongly accessible in other cell types. We speculate that these regions may display transient accessibility due to the intrinsic tendency of their underlying DNA sequences to disfavor nucleosome positioning [26-29].

While some accessible regions are stabilized at certain degrees by transcription factors, chromatin remodelers, and histone posttranslational modifications in specific cell types, others remain only transiently accessible. Nevertheless, these transiently accessible regions can still be captured by Tn5 transposase in a small number of cells. The extent to which these CREs that are accessible in a very small number of cells within a theoretically homogeneous population are important for the function of the cell and the regulation of transcription remains unknown. We have no doubt that CREs with a higher probability of accessibility would play a more significant role in transcriptional regulation. However, we cannot rule out the possibility that low-probability accessibility is required for a rapid response to signaling pathways induction. This is consistent with our results, where TGFβ increased the probability of accessibility for thousands of already accessible CREs, many of which were accessible in only a few cells (out of 3,925) before TGFβ treatment.

We have shown that *Pa*_*x*_ and Δ*Pa*_*x*_ provide information complementary to MACS2 scores and log_2_(FC) of the snATAC-seq signal. They offer a more realistic representation of accessibility within the cell population, while avoiding the ambiguity of whether an AR truly exists, since each region is described probabilistically as the likelihood of being accessible in the population, in a specific experiment.

Although the present study was performed using a single cell type, the approach can, in principle, be extended to more heterogeneous datasets, such as those derived from complex tissues comprising multiple cell types. In such cases, *Pa*_*x*_ could be calculated independently for each cell type after clustering. In future developments, the probabilistic framework described here could be extended to compare distinct cell types based on their full distributions of *Pa*_*x*_. Such comparisons would allow quantifying global and locus-specific differences in chromatin accessibility between cell populations, providing a more continuous and statistically grounded view of regulatory variation. This approach would enable the identification of subtle regulatory shifts that may not be captured by traditional threshold-based or fold-change analyses, offering a powerful direction for future studies of cell-type–specific chromatin landscapes.

Therefore, we think that this quantitative and probabilistic perspective on chromatin accessibility would enhance our understanding of the function of regulatory elements.

## METHODS

### Sample preparation and nuclei isolation

Normal Murine Mammary Gland (NMuMG) cells were cultured in DMEM supplemented with 10% fetal bovine serum (FBS; Cytiva-HyClone), 10 μg/ml insulin (Sigma), 100 µg/ml penicillin (Sigma), and 100 µg/ml streptomycin (Sigma). Cells were treated with either 5 ng/ml of vehicle solution (1 mg/ml BSA and 4 mM HCl) (control) or 5 ng/ml of TGFβ for 24 hours before trypsinization. Nuclei isolation was performed as described in 10X Genomics protocol for a final concentration of 5000-7000 nuclei/µl per sample. Cell lysis was performed by resuspending the cells in 200 µl of chilled Lysis Buffer and mixing by pipetting 10 times. The suspension was incubated in ice for 1 minute, after which 0.9 ml of chilled Wash Buffer was added to the lysate and mixed by pipetting 5 times. Cells were washed three times with 0.9 ml of Wash Buffer, pipetting 5 times, and centrifugated at 300 x g at 4ºC for 5 minutes. The resulting pellet was resuspended in 40 µl of chilled Nuclei Diluted Buffer. Approximately 50% of the nuclei were lost during cell lysis. 5000-7000 nuclei/µl were counted using a Neubauer chamber with Trypan Blue (Sigma), determining that more than 95% of cells were lysed (dead). Nuclei were diluted again to get the same nuclei concentration between samples. Nuclei integrity was tested by microscopy.

Isolated nuclei were then sent to CABIMER’s Genomic Unit to continue with the protocol. Joint scRNA- and snATAC-seq libraries were prepared following the 10X Genomics Chromium Next GEM Single Cell Multiome ATAC + Gene Expression user guide (CG000338). Briefly, 5 µl of each sample was used for nuclei isolation in the *Chromium controller* (10X Genomics), where nuclei were embedded inside lipidic droplets. Inside these droplets, tagmentation, RNA reverse transcription, barcode labeling and pre-amplification took place. Following, single-cell ATAC and RNA libraries were constructed, quantified using Qubit, and evaluated using a Bioanalyzer 2100 (High Sensitive DNA kit). Control and TGFβ24h single-cell Multiome libraries were sequenced with paired-end 100 bp reads on an Illumina NovaSeq 6000 instrument.

### Data pre-processing

Demultiplexing, genome alignment, and feature-barcode matrix generation were performed using the 10x Genomics cellranger-arc function (v2.0.2), using the mm10 reference genome (v2020-A). The output data were processed using Seurat (v4.3.0.1) [21] and Signac (v1.11.0) [22] R packages for scRNA-seq and snATAC-seq, respectively. Quality control was performed separately for scRNA-seq and snATAC-seq for each sample. Cells with between 1,000 and 100,000 ATAC fragments and a nucleosomal signal of at least 1.5 were kept. Cells with fewer than 1,000 and more than 40,000 reads for scRNA-seq, as well as cells with low TSS enrichment (< 1), were filtered out. Thus, 9,180 nuclei were retained from the initial 9,979 nuclei in the control condition. In the TGFβ24h condition, 4,068 nuclei were used for downstream analysis from the initial 4,201 nuclei. The TGFβ24h dataset was used for the differential analysis (see below).

For the control condition, analysis was performed either with 9,180 nuclei or with 3,925 nuclei. To obtain the 3,925 dataset, we randomly downsampled the control condition. Next, ATAC fragments were filtered to include only barcodes from the 3,925 nuclei using a streaming hash-based lookup, ensuring efficient selection of cell-specific reads. Thus, from the 229,796,725 fragments in the 9,180 nuclei, we retained 86,262,055 fragments in the 3,925 nuclei.

Peak calling was then performed using MACS2 (v2.2.7.1) [41] with --gsize 1.87e9 and --pvalue 0.1, resulting in 236,128 peaks for the 3,925 nuclei and 299,640 peaks for the 9,180 nuclei.

In the case of DMSO-treated LNCaP cells from [30] (GSE168667), snATAC-seq data were analyzed using Signac R package with the hg38 annotation for chromatin assay creation. Cells with 100-20,000 ATAC counts, a nucleosome signal lower than 10, and TSS enrichment greater than 0.5 were kept for downstream analysis. Thus, after filtering, analysis was performed with 4,269 nuclei, from the initial 4,436 nuclei. Peak calling was performed using MACS2 with --gsize 2.7e9 and --pvalue 0.1, resulting in 368,265 accessible regions (ARs).

### snATAC-seq fragments matrix processing

A snATAC fragments matrix for the control condition with n = 3,925 cells, containing the number of fragments per peak for each cell, was constructed (matrix dimensions: 236,128 ARs x 3,925 cells). Then, the total number of fragments per AR, the number of positive cells for each AR (cells in which an AR is detected with at least one fragment), and the number of cells in which ARs contain at least two fragments, were computed from this matrix. For all detected ARs, the average number of fragments per positive cell was also computed (see https://github.com/elesanesc/scPeakProbability.git). A control condition fragment matrix with 9,180 cells (299,640 ARs x 9,180 cells), and a matrix for the LNCaP experiment (368,265 ARs x 4,269 cells) were also obtained, with parameters computed as described above. Frequency distributions of the number of positive cells per AR were determined using standard methods in R (see https://github.com/elesanesc/scPeakProbability.git).

### Peak overlapping between different datasets

Overlapping peaks between Encode open region list (https://genome.ucsc.edu/cgi-bin/hgTrackUi?db=mm10&g=encodeCcreCombined), or Zu et al. [24], and our data were determined using findOverlapsOfPeaks() function from the ChIPpeakAnno R package (v3.34.1). Overlapping peaks between the peak calling results from the 3,925- and 9,180-nuclei datasets were determined in the same way.

### Differential snATAC-seq analysis between TGFβ-treated and control cells

To maintain comparable sequencing depth across conditions, TGFβ fragment counts were probabilistically downsampled to match the total fragment number of the subsampled control condition using a streaming Bernoulli sampling approach (retention probability *p = N_vehicle / N_TGFβ* ≈ *0*.*663;* fixed random seed = 1234). This resulted in 86,198,972 fragments.

Peak calling for the TGFβ condition was then performed using MACS2 (v2.2.7.1) [41] as previously described (with --gsize 1.87e9 and --pvalue 0.1 options), resulting in 290,875 peaks for the 4,068 nuclei. Next, a list of consensus peaks from the 3,925 nuclei in the control condition and the 4,068 nuclei in the TGFβ24h condition was generated using the reduce() function from the GenomicRanges R package (v1.52.1) [43], resulting in 350,123 ARs. Fragments were quantified in each AR for each condition using the FeatureMatrix() function from Signac, and the resulting datasets were then merged into a single Seurat object for downstream processing. snATAC-seq data were then processed using FinTopFeatures() with min.cutoff = 5, RunTFIDF(), and RunSVD() functions from the Signac package. Differential ARs analysis between the 350,123-consensus peak list from TGFβ24h and control conditions from the snATAC-seq datasets was performed using the FindMarkers() function from the Seurat package, with the parameters assay = “peaks”, min.pct = 0, test.use = “wilcox”, min.cells.feature = 1, and min.cells.group = 1. ARs with a |FC| > 0.5 and a p-value < 0.05 were considered statistically different between the two conditions. 111,359 ARs displayed increased accessibility after TGFβ treatment, while 64,077 showed decreased accessibility. Gene ontology (GO) functional categories were analyzed using David [42], and gene distributions were analyzed using ShinyGO [43].

### snATAC-seq count matrix processing for differential peaks

Fragment count matrices for the ARs with increased or decreased accessibility were extracted and computed as described above. The resulting matrices had the following dimensions: 111,359 ARs x 3,925 cells (control matrix for ARs with increased accessibility after TGFβ treatment), 64,077 ARs x 3,925 cells (control matrix for ARs with decreased accessibility after TGFβ treatment), 111,359 ARs x 4,068 cells (TGFβ matrix for ARs with increased accessibility after TGFβ treatment), and 64,077 ARs x 4,068 cells (TGFβ matrix for ARs with decreased accessibility after TGFβ treatment). Next, parameters were calculated as described above.

### scRNA-seq data analysis and UMAPs

scRNA-seq data were normalized using SCTtransform() function from the Seurat package. Neighbours were identified, and a UMAP projection was performed using 2:50 principal components from scRNA-seq’s PCA reduction and 2:50 principal components from snATAC-seq’s LSI (latent semantic indexing) reduction.

UMAPs showing gene expression levels and ATAC signal were plotted using the FeaturePlot_scCustom() function from the R package scCustomize (v3.0.1; Marsh SE (2021). scCustomize: Custom Visualizations & Functions for Streamlined Analyses of Single Cell Sequencing. https://doi.org/10.5281/zenodo.5706430. RRID:SCR_024675.)

### Poisson approximation to the “ball into bins” probability

The problem involves *m* balls that are thrown independently into *n* bins of identical size. Let *L* be a discrete random variable counting the number of balls into a bin. For n ≈ m, *L* can be approximated using the Poisson distribution. Then, the probability mass function for *L* is

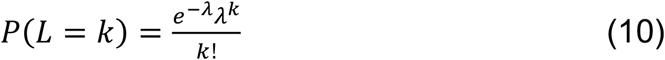

where λ is the mean (*m/n*) and *k* is the number of fragments at AR_*L*._ We verified that the mean-variance relationship of fragment counts was broadly consistent with a Poisson distribution in our datasets (Supplementary Figure S5).

Therefore, we calculate the probability of an empty bin:

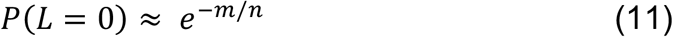

the probability of having at least one fragment is

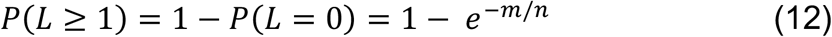

and the probability of having at least two fragments is

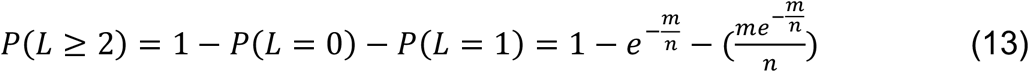

The corresponding expectancies, applying the linearity of the expectancy property, are

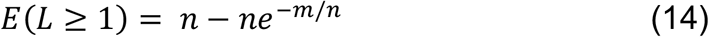

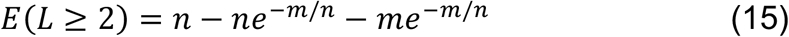

Equations 14 and 15 are very similar to equations 5 and 6 and can be also used to calculate *Pa*_*x*_.

### “Balls into bins” simulation

Computational simulations were conducted based on the classical *balls into bins* problem, with the aim of analyzing the distribution of balls across bins under uniform random assignment. In this simulation, the total number of fragments per AR was treated as “balls” (*m*), and the number of cells was considered as “bins” (*n*). In each iteration of the experiment, a set of *n* bins was considered, and *m* balls were thrown, with each ball being independently assigned to a randomly selected bin. The assignment was performed with replacement, allowing multiple balls to fall into the same bin. The simulation was implemented in R, using the sample() function to perform the random allocation of balls. Specifically, in each trial, the command sample(1:n, m, replace = TRUE) was executed, generating a sequence of *m* integers representing the bin indices for each ball. The number of balls per bin was then counted using the table() function. This process was repeated a number of times corresponding to the number of AR we want simulate. The results from each repetition were stored in a matrix for subsequent statistical analysis.

### Probability of accessibility (*Pa*_*x*_) calculation

We implemented the *Pa*_*x*_ calculation with Equation 9 in the scPaX package (v0.1.0; GitHub: https://github.com/elesanesc/scPaX). Briefly, scPaX accepts a Seurat object containing a “peaks” assay (or any fragment-count matrix), filters out zero-signal regions, and then fits the nonlinear model (Equation 9) using the base R function nls() from the stats package to estimate the hyperparameter *a*. The package returns a data frame with fragment counts and observed versus fitted *Pa*_*x*_ values, and generates publication-ready visualizations (scatter of *Pax* vs. fragment count; histogram of log_10_(*Pax*) distribution) via ggplot2. This package provides a reproducible and user-friendly tool for comparative chromatin accessibility analyses across any snATAC-seq dataset.

### MAE, RMSE and rRMSE

To evaluate the models’ or the simulation proximity to the observed data we have used the mean absolute error (MAE), root mean square error (RMSE) and relative RMSE (rRMSE).

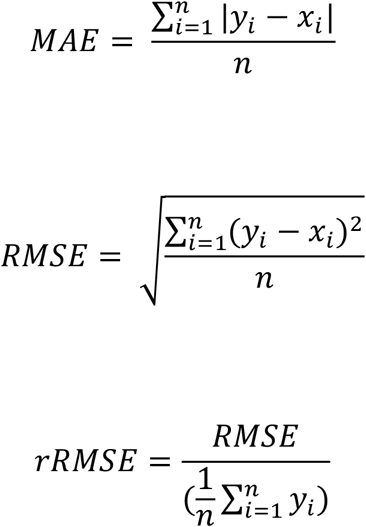

where *y*_*i*_ are the observed values and *x*_*i*_ are the predicted values.

### Other Statistical analysis

Statistical and graphical data analyses were performed using Prism 8 (Graphpad) software and Excel v15.24 (Microsoft). To determine the significance of the differences between groups, two-tailed Student’s t-tests with confidence interval of 95% were computed. Data were judged to be statistically significant when p value < 0.05 in applied statistical analyses. The coefficient of determination (R^2^) was also computed with Prism 8. The horizontal black line of the violin plots represents the median value, the 25th and 75th percentiles.

## Data and availability

The datasets supporting the conclusions of this article are available in the GEO repository with the accession code GSE285192 (Single-cell Multiome datasets generated in this study). The following secure token has been created to allow review of record GSE285192: gpklgscetvenzql. snATAC-seq data of DMSO-treated LNCaP cells from [33] were obtained from GSE168667 accession number. Bulk RNA-seq data from NMUMG cells [36] were obtained from GSE226954. ENCODE open region list was downloaded from UCSC Genome Browser (https://genome.ucsc.edu/), and the CREs list from adult mouse brain was extracted from Zu et al. [25]. Custom code and scripts used for analysis and for *Pa*_*x*_ determination are available at GitHub repository (https://github.com/elesanesc/scPeakProbability.git and https://github.com/elesanesc/scPaX).

## ACKNOWLEDGEMENTS

We thank J. A. Nepomuceno and Manuel Rodríguez-Paredes for critical reading of the manuscript. We thank Manuel Rodríguez-Paredes and Frank Lyko for introducing E.S.-E. to the computational analysis of snATAC-seq data. We thank E. Andújar and M. Pérez from the CABIMER Genomics Unit for NGS. Research in the J.C.R. lab was funded by the Spanish Ministry of Science and Innovation MCIU/AEI/10.13039/501100011033/ (grant number PID2023-149538NB-I00), by the regional Government of Andalusia (“Andalucía Biotec Salud”, BIOT22_00018_2.) and the European Union FEDER “A way to build Europe” program. CABIMER is a Center partially funded by the Junta de Andalucía. E.S.-E. is recipient of FPU fellowships from the Spanish Ministry of Education.

## AUTHOR CONTRIBUTIONS

Conceptualization: J.C.R. Methodology: E.S.-E. and J.C.R. Investigation: E.S.-E. and J.C.R. Data analysis: E.S.-E. and J.C.R. Visualization: E.S.-E. and J.C.R. Project administration, J.C.R.; Funding acquisition, J.C.R. Supervision, J.C.R. Review of the manuscript: D.R. Writing: E.S.-E. and J.C.R.

## INCLUSION AND DIVERSITY

We support inclusive, diverse, and equitable conduct of research.

