## Supplementary Figures for "Toward a probabilistic definition of chromatin accessible regions at the single-cell level"

Figure S1. Characterization of snATAC-seq data and detected accessible regions (AR)
Figure S2. Supplementary figures of the “balls into bins” model and the Probability of accessibility data.
Figure S3. Characterization of $Pa_x$ values.
Figure S4. Effect of TGF$\beta$ on accessibility.
Figure S5. Relationship between the variance and the mean of fragment counts across cells.

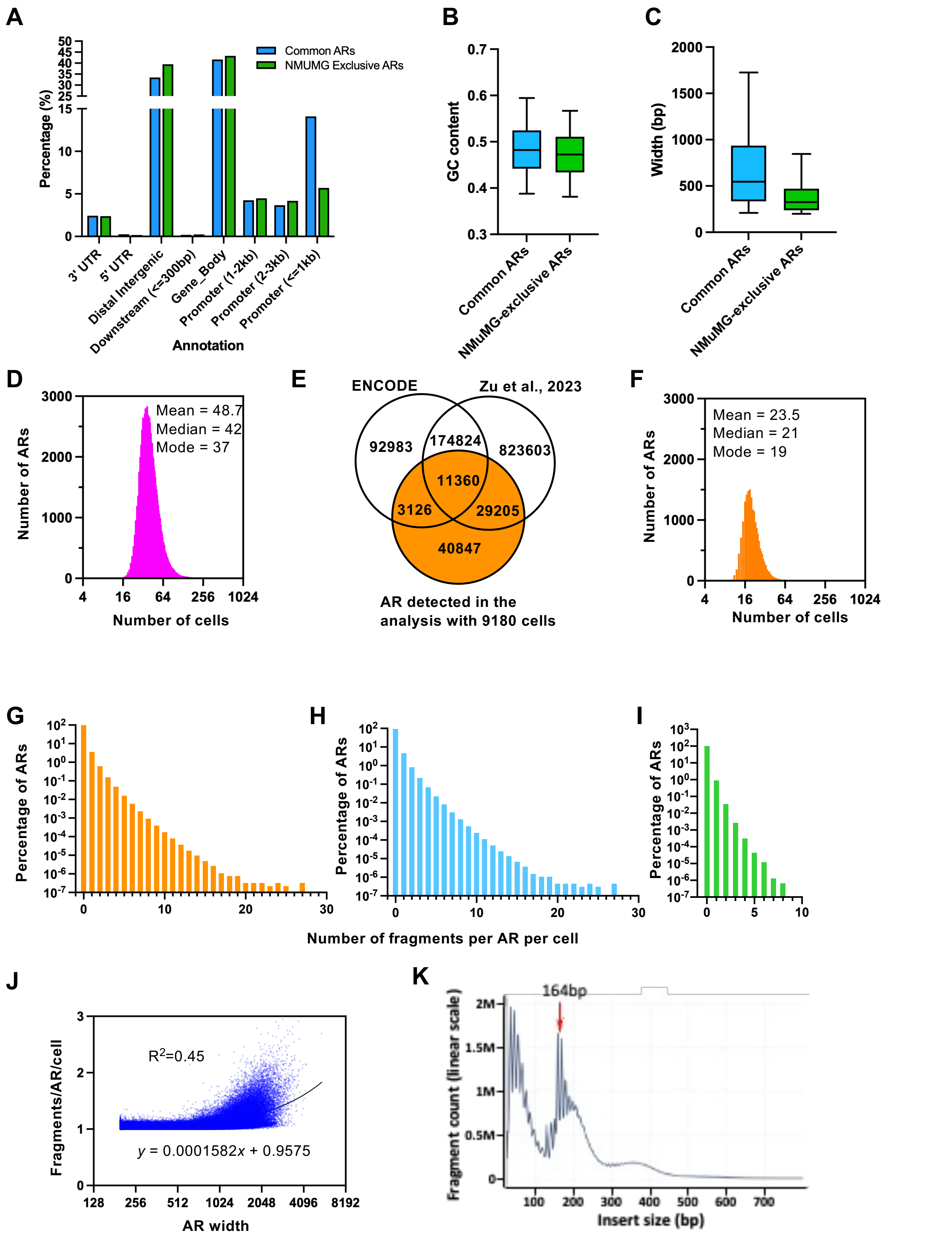


Supplementary Figure 1

**Figure S1. Characterization of snATAC-seq data and detected accessible regions (AR). A.** Type of regulatory elements present in common and NMUMG-exclusive ARs. **B.** GC content of common and NMUMG-exclusive ARs. **C.** Width of common and NMUMG-exclusive ARs. **D**. Frequency distributions of the number of positive cells for each AR detected exclusively in the analysis with 9180 cells. **E.** Overlapping between detected exclusively in the analysis with 9180 cells and mouse ARs previously described at the ENCODE mouse Registry of Candidate cis-Regulatory Element [23] and the candidate CREs from mouse adult brain [24]. **F.** Frequency distributions of the number of positive cells for each AR detected exclusively in the analysis with 3925 cells. **D-F**. Please note that the x-axis is in $log_2$ scale. **G-I.** Distribution of the number of fragments per AR per cell for all ARs detected in this study (G), for the common ARs (H) or for the NMUMG exclusive ARs (I). In contrast to Figure 1H, here all cells (not only positive cells) are taken into account. Please note that the y-axis is in $log_{10}$ scale **J.** Correlation between the number of fragments per AR per positive cell and the width of the AR. Coefficient of determination ($R^2$) of the linear regression is provided. Please note that the x-axis is in $log_2$ scale. **K.** snATAC-seq fragments distribution.

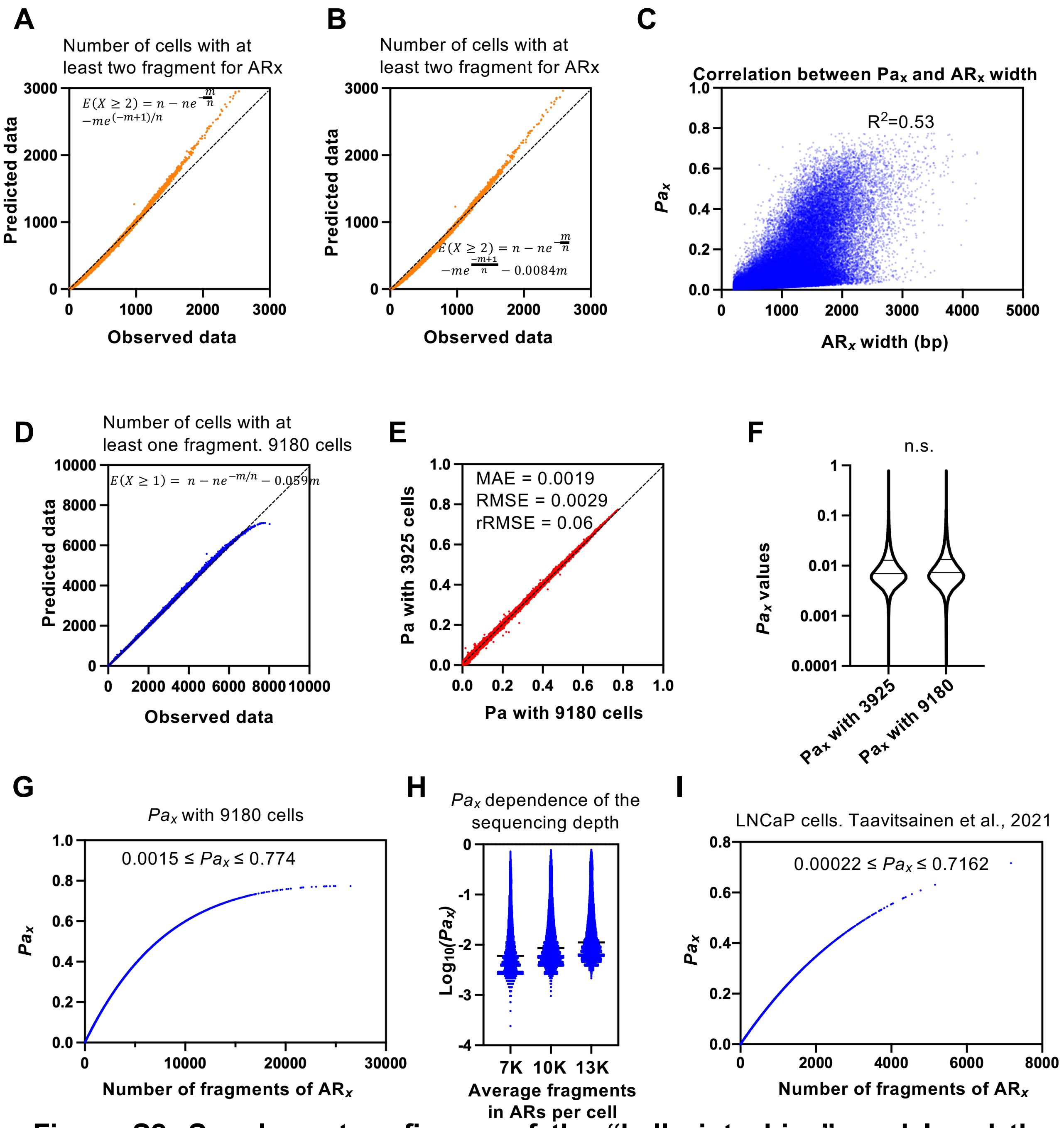


**Figure S2. Supplementary figures of the "balls into bins" model and the Probability of accessibility data. A, B.** Observed versus Predicted plots. Predicted values were estimated using the indicated equation were $n$ = 3925 and $m$ is the total number of fragments for every $AR_x$. **C.** Scatter plot showing the correlation between $Pa_x$ and the width of $AR_x$. Coefficient of determination ($R^2$) of the linear regression is provided. **D** Observed versus Predicted plots. Predicted values were estimated using the indicated equation were $n$ = 9180 and $m$ is the total number of fragments for every $AR_x$. **E.** Comparison of $Pa_x$ values computed using 3925 or 9180 cells. Mean absolute error (MAE), root mean square error (RMSE) and relative RMSE (rRMSE). **F.** Violin plots showing the distribution of $Pa_x$ values computed using 3925 or 9180 cells. **G.** Scatter plot showing how $Pa_x$ depends on the total number of fragments detected for $AR_x$. $Pa_x$ were computed using 9180 cells. **H.** Comparison of $Pa_x$ values computed using 3925 cells and the indicated average number of AR fragments per cell. The different numbers of average AR fragments per cell were obtained by downsampling. **I.** Scatter plot showing how $Pa_x$ depends on the total number of fragments detected for $AR_x$ from LNCaP cells, using data from reference (30).

Supplementary Figure 2

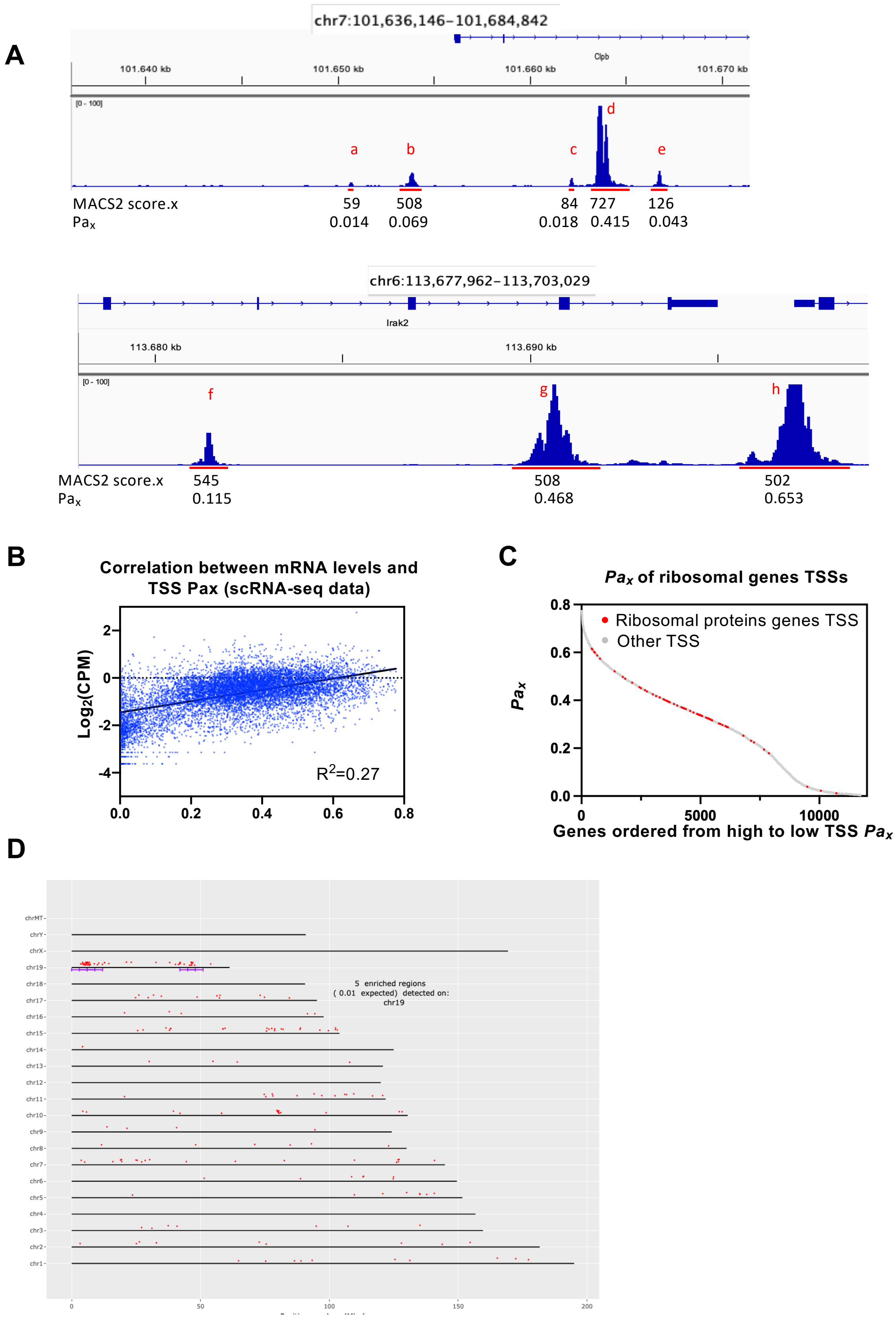


**Figure S3. Characterization of *Pa$_x$* values. A.** Genome browser snapshot of two genomic regions showing pseudobulk snATAC-seq signal of NMUMG cells under control conditions. *Pa$_x$* and *MACS2 score.x* values of the indicated ARs are shown. ARs were designed with letters in red. **B.** Correlation between *Pa$_x$* of TSS and mRNA levels of the corresponding genes using scRNA-seq data of the scMULTIOME experiment. **C.** Top-down ordered *Pa$_x$* values of TSS of all expressed genes. Ribosomal genes are represented by red dops. **D**. Genomic distribution of the genes with the top 200 TSS *Pa$_x$*.

Supplementary Figure 3

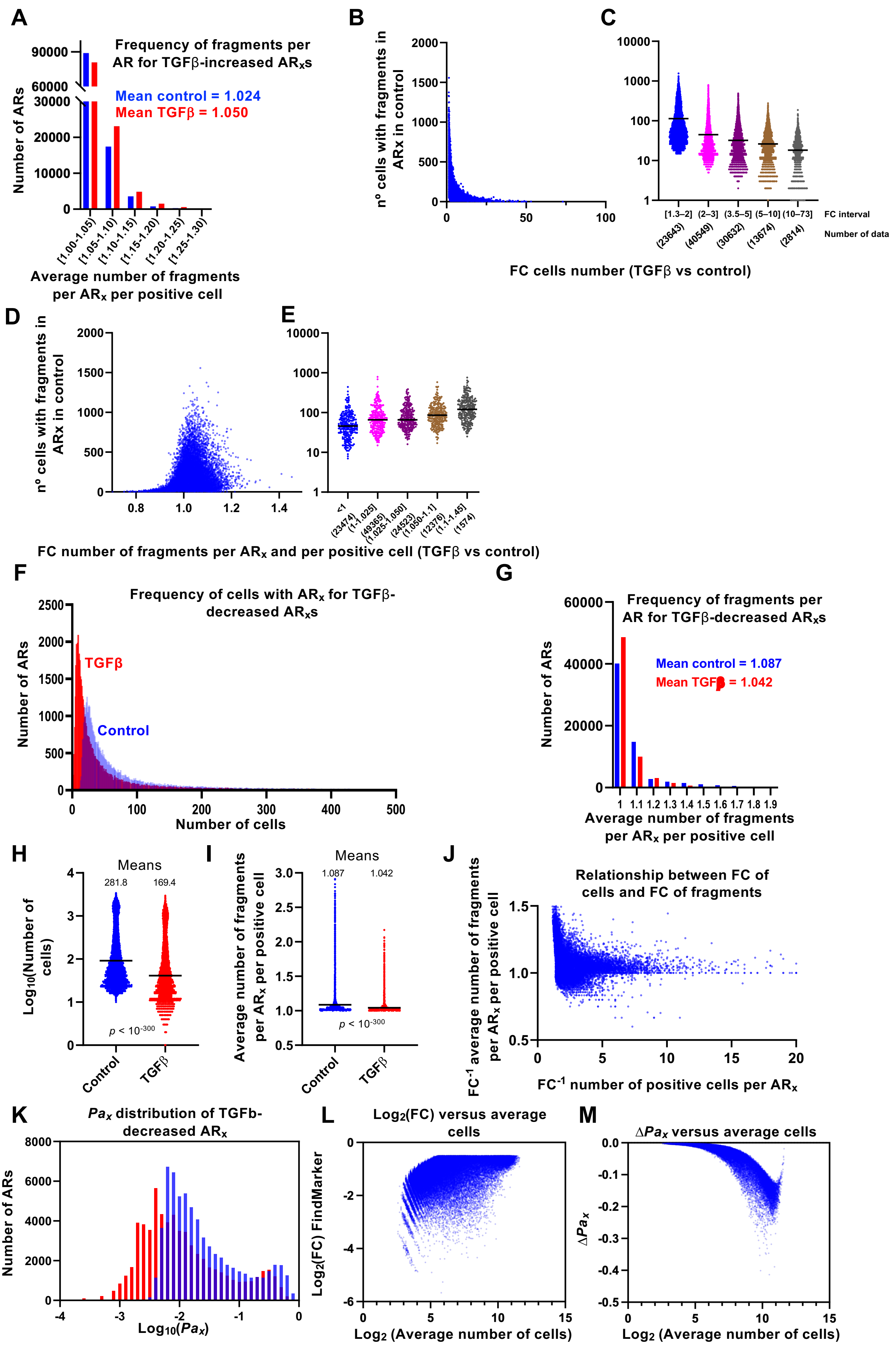


Supplementary Figure 4

**Figure S4. Effect of TGFβ on accessibility. A.** Frequency distributions of the average number of fragments per ARx and per positive cell for TGFβ-increased $AR_x$s. **B.** Relationship between the fold change (FC) (TGFβ vs control) of the number of positive cells of TGFβ-increased $AR_x$s and the total number of positive cells in the control condition. **C**. Quantification of B. FCs were binned into 5 intervals and the values of the number of positive cells were potted in violin plots. **D.** Relationship between the fold change (FC) (TGFβ vs control) of the average number of fragments per ARx and per positive cell, for TGFβ-increased $AR_x$s, and the total number of positive cells in the control condition. **E**. Quantification of D. FCs were binned into 5 intervals and the values of the number of positive cells were potted in violin plots. **F-M.** Data displayed in this figures correspond to the 64,077 $AR_x$s that decreased accessibility upon the TGFβ treatment (TGFβ-decreased $AR_x$s). Data for TGFβ-incresed $AR_x$s are shown in Figure 4A to 4J. **F.** Frequency distributions of the number of positive cells for the TGFβ-decreased $AR_x$s. Only the first part of the distribution (1-500 cells) is shown. **G.** Frequency distributions of the average number of fragments per $AR_x$ and per positive cell. **H.** Distribution of the number of positive cells in TGFβ-treated and non-treated cells. **I.** Distribution of the average number of fragments per $AR_x$ per positive cell in TGFβ-treated and non-treated cells. **J.** Relationship between the inverse of the fold change ($FC^{-1}$, TGFβ versus control) of the number of positive cells for $AR_x$ and the $FC^{-1}$ of the average number of fragments per $AR_x$ per positive cell. **K**. Comparison of the distributions of frequencies of $Pa_x$ under control conditions and after TGFβ addition, for all TGFβ-decreased $AR_x$s. Data are binned into intervals of 0.1 units. **L.** MA plot showing $Log_2FC$ (from FindMarkers(), Seurat) values versus $Log_2$(average number of cells) for TGFβ-decreased $AR_x$. **M**. MA plot showing $\Delta Pa_x$ values versus $Log_2$(average number of cells) for TGFβ-decreased $AR_x$.

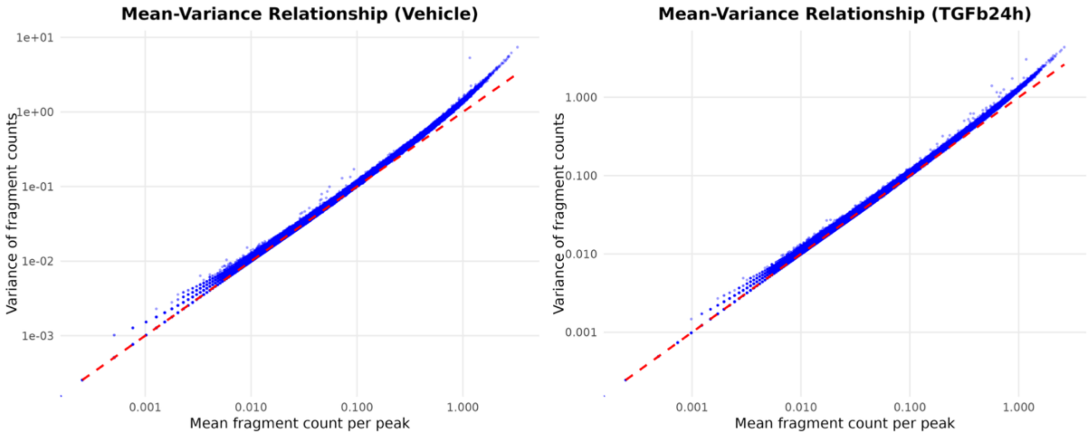


**Figure S5. Relationship between the variance and the mean of fragment counts across cells.** Each dot represents one AR region. The variance of fragment counts is approximately equal to the mean, consistent with a Poisson distribution (red line).
